# Convergent, functionally independent signaling by mu and delta opioid receptors in hippocampal parvalbumin interneurons

**DOI:** 10.1101/2021.04.23.441199

**Authors:** Xinyi Jenny He, Janki Patel, Connor E. Weiss, Xiang Ma, Brenda L. Bloodgood, Matthew R. Banghart

## Abstract

Functional interactions between G protein-coupled receptors are poised to enhance neuronal sensitivity to neuromodulators and therapeutic drugs. Mu and Delta opioid receptors (MORs and DORs) can interact when overexpressed in the same cells, but whether co-expression of endogenous MORs and DORs in neurons leads to functional interactions is unclear. Here, we show that both MORs and DORs inhibit parvalbumin-expressing basket cells (PV-BCs) in hippocampal CA1 through partially occlusive signaling pathways that terminate on somato-dendritic potassium channels and presynaptic calcium channels. Using photoactivatable opioid neuropeptides, we find that DORs dominate the response to enkephalin in terms of both ligand-sensitivity and kinetics, which may be due to relatively low expression levels of MOR. Opioid-activated potassium channels do not show heterologous desensitization, indicating that MORs and DORs signal independently. In a direct test for heteromeric functional interactions, the DOR antagonist TIPP-Psi does not alter the kinetics or potency of either the potassium channel or synaptic responses to photorelease of the MOR agonist DAMGO. Thus, despite largely redundant and convergent signaling, MORs and DORs do not functionally interact in PV-BCs. These findings imply that crosstalk between MORs and DORs, either in the form of physical interactions or synergistic intracellular signaling, is not a preordained outcome of co-expression in neurons.

## Introduction

G protein-coupled receptors (GPCRs) regulate cellular physiology through a diverse but limited number of intracellular signaling pathways. In neurons, signaling through multiple GPCRs expressed in the same cell can converge on the same molecular effectors (*e.g.* ion channels) to regulate neurophysiological properties such as cellular excitability and neurotransmitter release. Although GPCRs that engage the same family of G proteins (Gα_s_, Gα_i/o_ or Gα_q_) are poised to functionally interact through convergent biochemical signaling, it is not clear *a priori* whether such interactions would actually occur. Examples of interactions include functional synergy, when activation of one receptor subtype enhances activity at the other, or reciprocal occlusion, when the receptor subtypes compete for the same pool of effector molecules. Alternatively, GPCRs have been proposed to functionally interact through the formation of receptor heteromers, such that conformational changes due to ligand binding at one receptor shape agonist-driven signaling at the other.

Mu and delta opioid receptors (MORs and DORs) are both Gα_i/o_-coupled GPCRs that are activated by endogenous opioid neuropeptides such as enkephalin to suppress neuronal excitability and synaptic output. MORs are the primary target of widely used opiate analgesics (*e.g.* morphine, fentanyl) that are plagued by tolerance, high potential for addiction, and a propensity to cause respiratory depression. MORs and DORs have been proposed to functionally interact such that DOR-targeting drugs could reduce the clinical liabilities of MOR-targeting analgesics. For example, either pharmacological suppression or genetic removal of DOR attenuates morphine tolerance (Abdelhamid *et al.*, 1991; Sánchez-Blázquez, García-España and Garzón, 1997; Zhu *et al.*, 1999). Furthermore, co-administration of MOR and DOR agonists produces spinal, supraspinal and peripheral analgesic synergy (Porreca *et al.*, 1987; Schuster *et al.*, 2015; Bruce *et al.*, 2019). In contrast, antagonism of one receptor has been reported to enhance agonist-driven activity at the other receptor in assays using heterologous receptor expression. These observations have been interpreted to support the existence of MOR/DOR heteromers that interact through direct allosteric coupling (Fujita, Gomes and Devi, 2015; Cahill and Ong, 2018). MOR/DOR heteromers have been specifically implicated as potential therapeutic targets for the treatment of pain, as intrathecal coadministration of the DOR-selective antagonist TIPP-Psi with morphine produces stronger analgesia than morphine alone (Gomes *et al.*, 2004). Due to the clinical potential of therapeutic approaches that simultaneously engage MORs and DORs, understanding the mechanisms that underlie their potential for functional interactions is of great importance.

Relatively few studies have investigated functional interactions between endogenous MORs and DORs using sensitive measurements of cellular physiology with the single-cell resolution required to implicate cell-autonomous interactions, as opposed to circuit-level effects. In recordings from neurons in the nucleus raphe magnus after upregulation of DORs in response to chronic morphine treatment, MORs and DORs were found to synergistically suppress inhibitory synaptic transmission through a PKA-dependent pathway, but evidence of heteromers was not observed (Zhang and Pan, 2010). Indicative of functionally independent signaling, a more recent study that assessed endogenous MOR and DOR trafficking in spinal dorsal horn neurons did not find evidence for co-internalization after intrathecal administration of either the DOR-selective agonist SNC80 or the MOR-selective agonist [D-Ala^2^, NMe-Phe4, Gly-ol5]enkephalin (DAMGO) (Wang *et al.*, 2018). In contrast, recordings from ventral tegmental area neurons suggested MOR/DOR interactions consistent with heteromer formation (Margolis *et al.*, 2017). In that study, TIPP-Psi enhanced DAMGO-evoked membrane potential hyperpolarization, and the MOR antagonist CTOP enhanced hyperpolarization evoked by the DOR agonists DPDPE and deltorphin II. However, at least some of the recordings were from dopamine neurons, which have been shown not to express *Oprm1* mRNA (Galaj *et al.*, 2020). Thus, in naïve mice, unequivocal evidence for functional interactions between endogenous MORs and DORs in the same neurons, and in particular, for the existence of MOR/DOR heteromers, is lacking.

In some brain regions, including the hippocampus, MORs and DORs are established to be co-expressed in the same neurons, such that the receptors and their downstream intracellular signaling pathways are poised to interact (Chieng, Christie and Osborne, 2006; Erbs *et al.*, 2015). In the hippocampus, activation of MORs in GABA neurons contributes to stress-induced memory deficits (Shi *et al.*, 2020), whereas DORs may contribute to spatial contextual cue-related memory retrieval (Le Merrer *et al.*, 2011, 2012, 2013). Recently, we reported that MORs and DORs both contribute to opioid-mediated suppression of perisomatic inhibition in the CA1 region of hippocampus, consistent with previous studies of MOR and DOR modulation of synaptic transmission (Glickfeld, Atallah and Scanziani, 2008; Piskorowski and Chevaleyre, 2013; Banghart, He and Sabatini, 2018). In fact, MORs and DORs are well established to regulate inhibitory synaptic transmission in CA1 (Zieglgänsberger *et al.*, 1979; Nicoll, Alger and Jahr, 1980; Lupica and Dunwiddie, 1991; Lupica, Proctor and Dunwiddie, 1992; Lupica, 1995; Svoboda and Lupica, 1998; Svoboda, Adams and Lupica, 1999; Rezaï *et al.*, 2012). Although a substantial body of work indicates co-expression of MOR and DOR in CA1 parvalbumin basket cells (PV-BCs), which are a primary source of perisomatic inhibition (Stumm *et al.*, 2004; Erbs *et al.*, 2012; Faget *et al.*, 2012), a direct comparison of their neurophysiological actions has not been conducted.

In this study, we explored potential interactions between MORs and DORs in CA1 PV-expressing basket cells using recordings from hippocampal slices. In order to obtain precise and sensitive measures of receptor function, we optically probed native MORs and DORs using photoactivatable (caged) opioid neuropeptides (Banghart and Sabatini, 2012; Banghart, He and Sabatini, 2018). Using this approach, we found that MORs and DORs activate partially overlapping pools of somatodendritic potassium channels in PV-BCs, and suppress synaptic output from PV-BCs in a mutually occlusive manner. Despite their co-expression and functional redundancy, we did not find evidence of synergy or for heteromers, indicating that MOR and DOR signal in a parallel, functionally independent manner in PV-BCs.

## Results

### Occlusive suppression of hippocampal perisomatic inhibition by MORs and DORs

We first confirmed that both MORs and DORs are co-expressed in PV-BCs using fluorescence in-situ hybridization, which revealed that 78% (171/218) of *Parv* mRNA-containing neurons with cell bodies in and around stratum pyramidale contain both *Oprm1* and *Oprd1* mRNA (**Supporting Figure S1A, B**). To determine if both MORs and DORs are functional in PV-BCs, we virally expressed the light-gated cation channel Chronos in a Cre recombinase-dependent manner in the CA1 region of PV-Cre mice and measured the effects of the selective MOR and DOR agonists DAMGO and SNC162, respectively, on light-evoked synaptic transmission using electrophysiological recordings from pyramidal cells (PCs) in acute hippocampal slices (Klapoetke *et al.*, 2014). We chose SNC162 due to its exceptional selectivity for DOR over MOR (Knapp *et al.*, 1996). To maximize the relative contribution of perisomatic inhibition from PV+ basket cells, as opposed to dendrite-targeting PV+ bistratified cells, we restricted the area of illumination to a small region of stratum pyramidale around the recorded PC (**Figure 1A**). Bath perfusion of either DAMGO (1 μM) or SNC162 (1 μM) strongly reduced the optically-evoked IPSC (oIPSC) to a similar degree (**Figure 1B-D**). Sequential drug application only slightly increased the degree of suppression compared to either drug alone (DAMGO: 0.69 ± 0.05, n = 9 cells; SNC162: 0.70 ± 0.05, n = 9 cells; both: 0.76 ± 0.03, n = 18 cells; no significant differences, Ordinary one-way ANOVA) (**Figure 1D, Supporting Figure S1F**). In both cases, application of pairs of optical stimuli (50 ms apart) revealed small increases in the paired-pulse ratio (PPR) in the presence of the opioid agonist, consistent with a presynaptic mechanism of action for the opioid receptor (BL: 0.48 ± 0.03; DAMGO: 0.65 ± 0.09; n = 8 pairs; p = 0.0078, Wilcoxon matched-pairs signed rank test; BL: 0.59 ± 0.05; SNC162: 0.72 ± 0.06; n = 7 pairs; p = 0.047, Wilcoxon matched-pairs signed rank test) (**Figure 1E**). Interestingly, with sustained application, the effects of DAMGO, but not SNC162, appeared to desensitize slightly. These results reveal that both MORs and DORs suppress the output of PV-BCs in a mutually occlusive manner.

**Figure 1.**
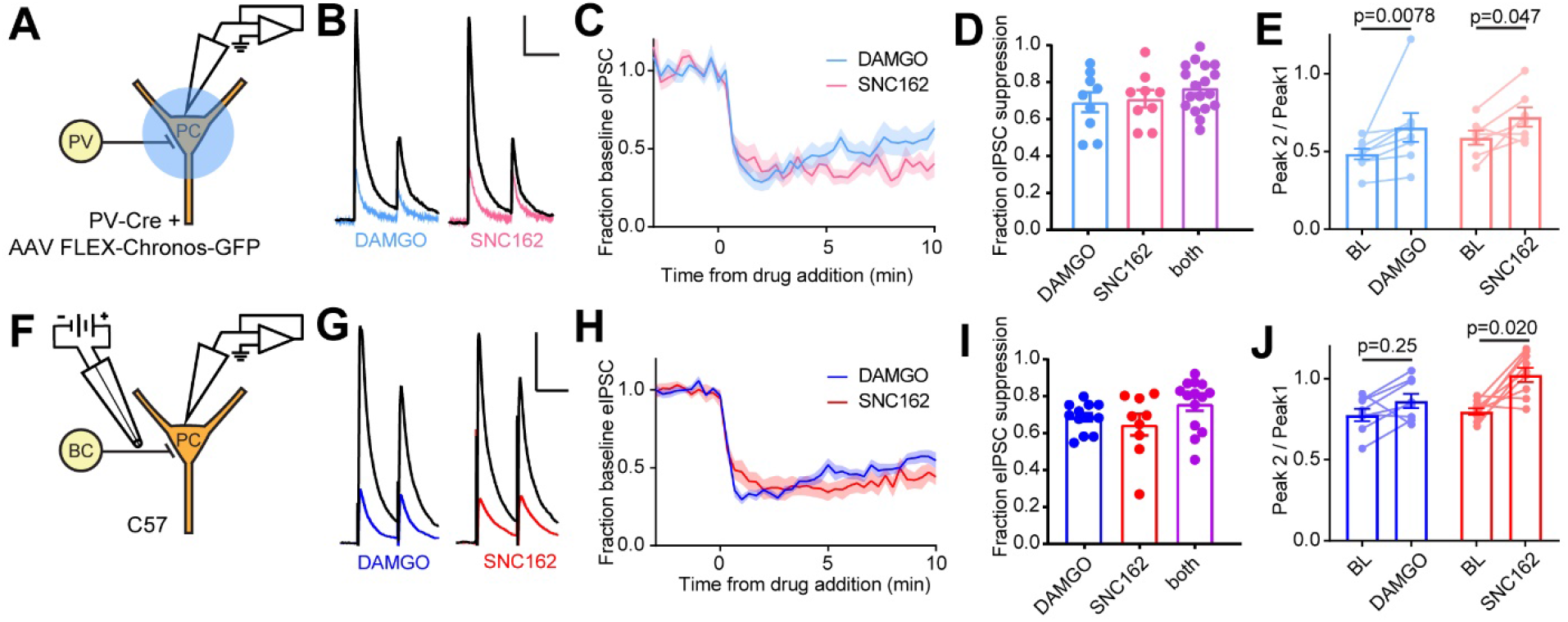
Electrophysiological recordings of opioid-sensitive synaptic output from hippocampal parvalbumin basket cells. **A.** Schematic of the experimental configuration for recording optogenetically-evoked inhibitory synaptic transmission in PV-Cre mice. **B.** Representative oIPSC pairs (50 ms interval) recorded from a pyramidal cell. Black traces are the average of 6 baseline sweeps, and colored traces are the average of 6 sweeps after addition of either DAMGO (1 uM, blue) or SNC162 (1 uM, red). Scale bars: x = 40 ms, y = 100 pA. **C.** Baseline-normalized, average oIPSC amplitude over time during bath application of DAMGO (n = 9 cells from 6 mice) or SNC162 (n = 9 cells from 7 mice). **D.** Summary data of double flow-in experiments, comparing oIPSC suppression by DAMGO or SNC162 alone, followed by the other drug. **E.** oIPSC paired pulse ratios (Peak 2/Peak 1), before (baseline, BL) and after drug addition. **F.** Schematic of the experimental configuration for recording electrically-evoked inhibitory synaptic transmission in wild type mice. **G.** Representative eIPSC pairs (50 ms interval) recorded from a pyramidal cell (as in **B**). Scale bars: x = 40 ms, y = 200 pA. **H.** Baseline-normalized, average eIPSC amplitude over time during bath application of DAMGO (n = 13 cells from 12 mice) or SNC162 (n = 9 cells from 5 mice). **I.** Summary data of double flow-in experiments with electrical stimulation (as in **D**). **J.** eIPSC paired pulse ratios (Peak 2/Peak 1), before and after drug addition.

To improve experimental throughput, we established an electrical stimulation protocol for preferential activation of PV-BC terminals by placing a small bipolar stimulating electrode in stratum pyramidale immediately adjacent to the recorded PC (**Figure 1F**). Recordings were made from PCs near stratum oriens, as these have been shown to receive BC input that is biased towards PV-BCs as opposed to CCK-BCs (Lee *et al.*, 2014). Consistent with only a minor contribution to the electrically evoked IPSC (eIPSC) from CB1R-expressing CCK-BCs, bath application of the CB1R agonist WIN55 (1 μM) resulted in only modest eIPSC suppression (0.25 ± 0.07, n = 8 cells), and application of DAMGO in the presence of WIN55 produced only slightly more suppression than DAMGO alone (DAMGO: 0.67 ± 0.02, n = 12 cells; WIN+DAMGO: 0.79 ± 0.03, n = 8 cells; p = 0.1072, Ordinary one-way ANOVA with Dunnett’s multiple comparisons test) (**Supporting Figure S1C-E**) (Glickfeld, Atallah and Scanziani, 2008). Under these electrical stimulation conditions, DAMGO and SNC162 again suppressed the eIPSC to a similar degree, with DAMGO producing slight desensitization, exhibited strong mutual occlusion (DAMGO: 0.67 ± 0.02, n = 12 cells; SNC162: 0.63 ± 0.06, n = 9 cells; both: 0.75 ± 0.04, n = 14 cells; no significant differences, Ordinary one-way ANOVA), and resulted in a small increase in PPR (BL: 0.78 ± 0.04; DAMGO: 0.86 ± 0.04; n = 8 pairs; p = 0.25, Wilcoxon matched-pairs signed rank test; BL: 0.80 ± 0.02; SNC162: 1.02 ± 0.04; n = 9 pairs; p = 0.02, Wilcoxon matched-pairs signed rank test) (**Figure 1 F-J, Supporting Figure S1F**). Thus, our electrical stimulation preferentially recruits opioid-sensitive PV-BCs, as the effects of DAMGO and SNC162 on the eIPSC and oIPSC were indistinct (no significant difference, Two-way ANOVA) (**Supporting Figure S1G)**.

MOR and DOR are thought to exhibit similar affinity for enkephalin, but how this translates to ligand efficacy at native receptors in neurons is not clear. In addition, receptor signaling kinetics could prove to be a sensitive means of detecting functional interactions. To compare the ligand sensitivity and receptor signaling kinetics of MORs and DORs, we turned to photoactivatable derivatives of the MOR and DOR agonist [Leu^5^]-enkephalin (LE) (**Figure 2A, top**) (Banghart and Sabatini, 2012). For quantitative pharmacology, we chose to use *N*-MNVOC-LE, which is highly inactive at both DOR and MOR (Banghart, He and Sabatini, 2018). In the presence of *N*-MNVOC-LE (6 μM), which is optimized for simultaneous activation of MORs and DORs, application of a strong 5 ms UV light flash 2 sec prior to an eIPSC produced a rapid, transient suppression of the IPSC that recovered within 1-2 minutes (**Figure 2A, B**). Varying UV light intensity in a graded fashion allowed us to rapidly obtain dose-response curves within a single recording. To assess the potency of LE at MORs and DORs, and the relative contributions of the receptors to the IPSC suppression by LE, we recorded dose-response curves in the absence and presence of the MOR- and DOR-selective antagonists CTOP (1 μM) and TIPP-Psi (1 μM), respectively (**Figure 2C**). We chose CTOP over its analog CTAP due to its higher selectivity for MORs. Whereas LE uncaging at the highest light power (84 mW) in the absence of opioid antagonists suppressed synaptic transmission by 63 ± 4%, activation of MORs or DORs alone, which were isolated by antagonizing with TIPP-Psi or CTOP, respectively, suppressed synaptic output by ~40% each. Although the extent of suppression achieved with caged LE was somewhat less than with bath application (**Figure 1I**), the relative contributions of MORs and DORs were similar in both experiments and consistent with mutual occlusion. The dose-response curve revealed that LE exhibits ~3-fold greater potency for DORs than MORs in regulating perisomatic inhibition (EC50 values in the absence (black, 3.28±0.47 mW) and presence of either CTOP (red, 2.29±0.61 mW) or TIPP-Psi (blue, 9.30±1.40 mW)). Moreover, DOR activation largely accounts for the actions of LE in the absence of antagonists. This could reflect greater affinity for DORs, or more efficacious signaling by DORs than MORs (**Figure 2D**).

**Figure 2.**
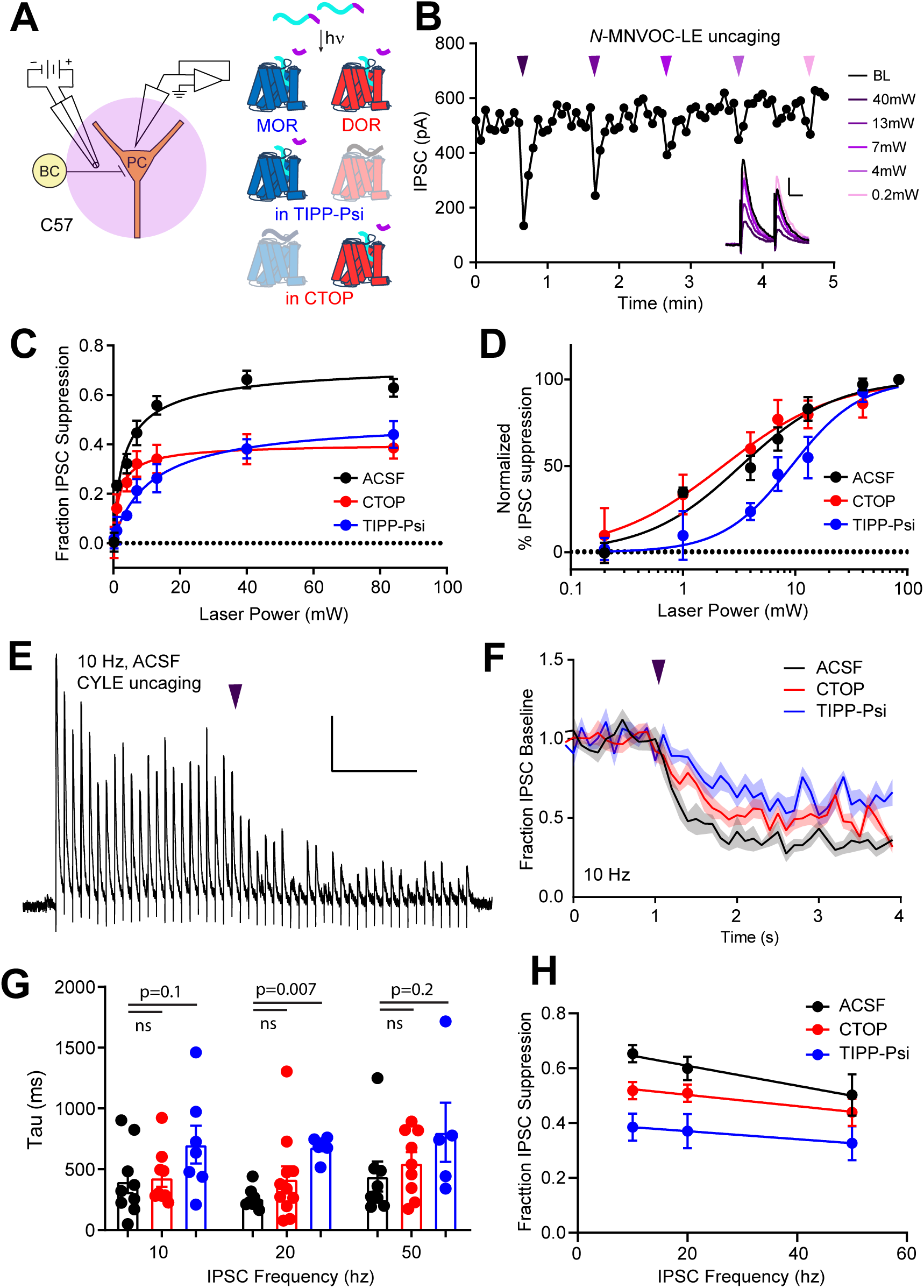
Characterization of the potency and kinetics of synaptic modulation by [Leu^5^]-enkephalin at mu and delta opioid receptors using caged peptides. **A. Left:** Schematic of the experimental configuration for photo-uncaging of opioid neuropeptides while recording electrically-evoked inhibitory synaptic transmission in wild type mice. **Right:** Schematic of photoreleasing [Leu^5^]-enkephalin (cyan) from *N*-MNVOC-LE or CYLE (cyan with purple caging group) in the presence of selective antagonists to isolate its action on either MOR (blue, in TIPP-Psi) or DOR (red, in CTOP). **B.** Example recording showing graded suppression of inhibitory synaptic transmission by uncaging *N*-MNVOC-LE at various light intensities. **Inset:** Example IPSCs before (black) and after LE uncaging at each light intensity. Scale bars: x = 20 ms, y = 100 pA **C.** Linear optical dose-response curves of IPSC suppression as a function of light intensity, in the absence (black, n = 6 -12 cells per laser intensity) and presence of either CTOP (red, n = 5-8 cells) or TIPP-Psi (blue, n = 4-10 cells). **D.** Logarithmic optical dose-response curves of the data in **C** normalized to the maximal IPSC suppression observed in each condition. **E.** Representative recording from a pyramidal cell demonstrating rapid suppression of IPSC amplitude in response to photoactivation of CYLE during 10 Hz trains of electrical stimuli. Purple arrow represents CYLE uncaging at 2 seconds into the 10 Hz train. Outward stimulus artifacts are removed for clarity. Scale bars: x = 1 sec, y = 100 pA **F.** Average, baseline subtracted and baseline-normalized IPSC amplitude showing the kinetics of synaptic suppression with electrical stimulation at 10 Hz in the absence (ACSF, n = 9 cells from 6 mice) and presence of either CTOP (n = 12 cells from 7 mice) or TIPP-Psi (n = 8 cells from 6 mice). **G.** Time constants of synaptic suppression in response to CYLE photoactivation with an 84 mW light flash at the indicated frequencies of synaptic stimulation. At 20 Hz, the time constant in TIPP-Psi was significantly greater than the time constant without any antagonists. **H.** Plot of IPSC suppression as a function of synaptic stimulation frequency.

We evaluated receptor signaling kinetics using the photoactivatable LE derivative CYLE, which photolyzes within tens of microseconds, such that receptor activation is rate-limiting (Banghart and Sabatini, 2012; Banghart, He and Sabatini, 2018). In order to sample synaptic transmission at frequencies sufficient to resolve receptor signaling kinetics, we drove eIPSCs in 5 s bouts at 10, 20 and 50 Hz, and photolyzed CYLE (6 μM) after synaptic depression had stabilized to a steady state (**Figure 2E**). To obtain the time-constants of synaptic suppression for each receptor, we repeated this experiment in the presence of the selective antagonists and fit the post-flash eIPSC amplitudes with a single exponential function (**Figure 2F**). The time constants we obtained for each pharmacological condition were similar for all three stimulus frequencies (**Figure 2G**). Whereas DOR (CTOP at 20 Hz, tau = 419 ± 105 ms, n = 11 cells) exhibited kinetics indistinct from the drug-free condition (ACSF at 20 Hz, tau = 259 ± 30 ms, n = 8 cells), the time-constant of MOR-mediated suppression was surprisingly slow (TIPP-Psi at 20 Hz, tau = 683 ± 36 ms, n = 6 cells; p = 0.007, Ordinary one-way ANOVA with Dunnett’s multiple comparisons test). We also observed that the extent of IPSC suppression correlated inversely with the frequency of synaptic stimulation, and that this was most pronounced in the absence of antagonists (**Figure 2H**).

Together, these results suggest that MOR and DOR suppress output from overlapping populations of PV-BC presynaptic terminals, and that this suppression is dominated by DOR, both in terms of sensitivity to LE and response kinetics.

### MORs and DORs suppress GABA release by inhibiting voltage-gated Ca^2+^ channels

At least two mechanisms of presynaptic inhibition by Gα_i/o_-coupled GPCRs have been established, but the pathways engaged by opioid receptors in PV-BCs are not known. One potential mechanism involves the inhibition of voltage-sensitive calcium channels (VSCCs) by Gβγ proteins (Bean, 1989), whereas the other involves direct suppression of SNARE proteins by Gβγ binding to the C-terminus of SNAP25 (Blackmer *et al.*, 2001; Gerachshenko *et al.*, 2005; Zurawski *et al.*, 2019; Hamm and Alford, 2020). The observed frequency-dependent synaptic suppression is consistent with both mechanisms, as Gβγ binding to VSCCs is reversed by strong depolarization, and elevated Ca^2+^ facilitates displacement of Gβγ from the SNARE complex by Ca^2+^-bound synaptotagmin (Park and Dunlap, 1998; Brody and Yue, 2000; Yoon *et al.*, 2007).

To ask if MOR and DOR inhibit presynaptic VSCCs in PV-BCs, we imaged action potential-induced Ca^2+^ transients in presynaptic boutons of PV-BCs using two-photon laser scanning microscopy. PV-BCs were targeted for whole cell current clamp recordings in PV-Cre; tdTom (Rosa-Lsl-td-Tomato (Ai14)) mice with the small molecule Ca^2+^ indicator Fluo5F included in the recording pipette (**Figure 3A**). Line scans across putative boutons were obtained while triggering either one or five action potentials, before and after bath application of DAMGO, SNC162 or both drugs together (**Figure 3B**).

**Figure 3.**
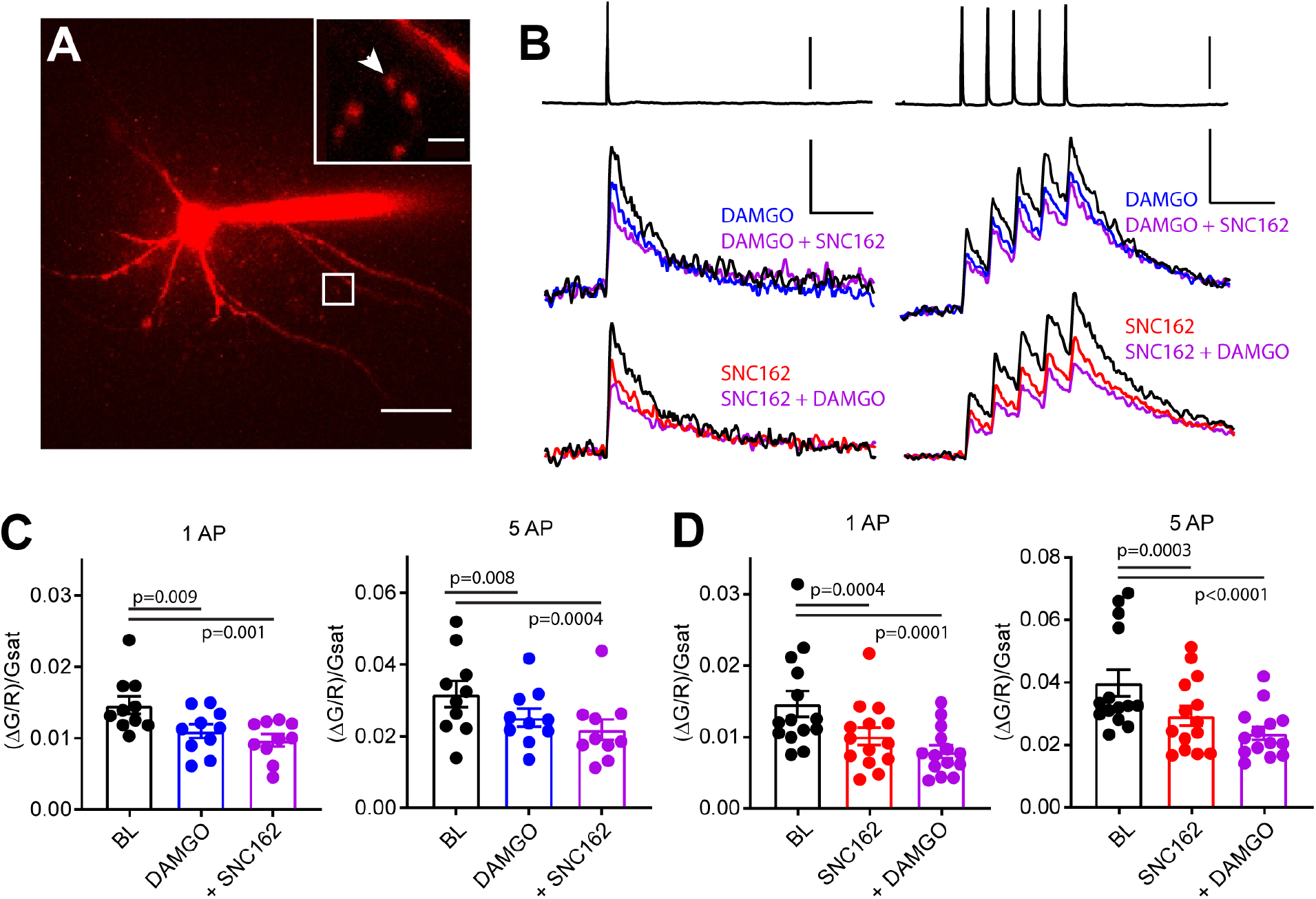
Axonal calcium imaging reveals that both mu and delta opioid receptors suppress presynaptic voltage-gated calcium channels. **A.** Two-photon image of a tdTomato-expressing basket cell filled with 30 μM Alexa 594 and 300 μM Fluo-5F in a brain slice taken from a PV-Cre; tdTom mouse. Scale bar: 50 μm. Inset shows the two axonal boutons where the line scan was carried out, with the orientation of the line scan indicated by the arrow. Scale bar: 5 μm. **B.** Example of either a single action potential (left) or five action potentials (right) triggered in the cell body (top), and the resulting averaged, presynaptic Ca^2+^ transients, before and after application of DAMGO (top, blue, n = 8 cells, 16 boutons), SNC162 (red bottom, n = 7 cells, 14 boutons), and both drugs (top and bottom, purple). The transients are measured as the change in green signal (ΔG) over red signal (R), divided by ΔG in saturating Ca^2+^ conditions (ΔG^sat^). Scale bars: top, 50 mV; bottom, x = 100 ms, y = 0.01 (left) or 0.02 (right) (ΔG/R)/Gsat. **C.** Summary of peak Ca^2+^ transients for DAMGO application in response to 1 AP (left) or 5 APs (right). 1 AP: BL 0.014 ± 0.001; DAMGO 0.011 ± 0.001; DAMGO+ SNC 0.010 ± 0.001 (p = 0.009 and p = 0.0001, n = 10 pairs, Repeated measures one-way ANOVA with Dunnett’s multiple comparisons test) 5AP: BL 0.032 ± 0.004; DAMGO 0.025 ± 0.002, DAMGO + SNC 0.022 ± 0.003 (p = 0.008 and p = 0.004, n = 10 pairs). **D.** Summary of peak Ca^2+^ transients for SNC162 application in response to 1 AP (left) or 5 APs (right). 1AP: BL 0.014 ± 0.002; SNC 0.010 ± 0.002; SNC + DAMGO 0.008 ± 0.001 (p = 0.0004 and p = 0.0001, n = 14 pairs, Repeated measures one-way ANOVA with Dunnett’s multiple comparisons test). 5AP: BL 0.039 ± 0.004; SNC 0.029 ± 0.003; SNC + DAMGO 0.023 ± 0.002 (p = 0.003 and p < 0.0001, n = 14 pairs)

Individually, DAMGO and SNC162 both caused a ~30% reduction in the peak ΔF/F evoked by either stimulation protocol (DAMGO 27.27% for 1 AP, 17.73% for 5 APs, SNC162 31.18% for 1 AP, 26.55% for 5 APs). When DAMGO and SNC162 were applied together, these presynaptic Ca^2+^ transients were suppressed by ~40%, on average (DAMGO then SNC162 40.95% for 1 AP, 38.92% for 5 APs, SNC then DAMGO 46.08% for 1 AP, 40.85% for 5 APs) (**Figure 3C, D**). Given the nonlinear Ca^2+^-dependence of vesicular fusion, a 30% reduction in presynaptic Ca^2+^ is consistent with the strong suppression of PV-BC IPSCs by MORs and DORs (Wu and Saggau, 1997). These results indicate that the inhibition of VSCCs by both MORs and DORs is the most likely mechanism accounting for their effects on inhibitory transmission. Furthermore, the marginal effect of adding a second drug suggests convergence on the same pool of VSCCs.

### Enkephalin generates large outward somato-dendritic currents in PV-BCs primarily through DORs rather than MORs

Gα_i/o_-coupled GPCRs, including both MORs and DORs, often hyperpolarize neurons by activating G protein-coupled inward rectifier K^+^ (GIRK) channels, as well as voltage-gated K+ channels, or by suppressing HCN channels (Williams, Egan and North, 1982; North *et al.*, 1987; Wimpey and Chavkin, 1991; Svoboda and Lupica, 1998). Although MORs were previously reported to activate outward currents in the somato-dendritic compartment of fast spiking CA1 BCs, the role of DORs has not been explored (Glickfeld, Atallah and Scanziani, 2008). To address this, we performed voltage-clamp recordings of opioid-evoked currents in tdTom-labeled cells in PV-Cre; tdTom mice (**Figure 4A**). At a holding potential of −55 mV, *N*-MNVOC-LE photoactivation using strong (84 mW) light flashes applied to the soma and proximal dendrites of the recorded neuron evoked rapidly rising outward currents that decayed over ~1 min, similar to previous observations in locus coeruleus (**Figure 4B, C**) (Banghart and Sabatini, 2012). Surprisingly, blocking MORs with CTOP had no measurable effect on the light-evoked current (ACSF: 81.7 ± 9.6 pA, n = 9 cells; CTOP: 82.5 ± 12.8 pA, n = 10 cells; not significant). In contrast, blocking DOR with TIPP-Psi greatly reduced the current amplitude (TIPP-Psi: 26.4 ± 4.8 pA, n = 11 cells; p = 0.006), and addition of both drugs completely abolished it (CTOP + TIPP-Psi: 7.1 ± 0.09 pA, n = 5 cells; p = 0.0009 and p = 0.009; Welch’s ANOVA with Dunnet’s T3 multiple comparisons test). Optical dose-response curves in the presence of each antagonist revealed a larger DOR-mediated than MOR-mediated current (**Figure 4D**). Similar to our observations with presynaptic receptors, LE exhibited greater potency at DORs than MORs in generating outward currents (EC50 values of ACSF: 17.55 ± 2.98 mW, CTOP: 7.59 ± 1.26 mW, TIPP-Psi: 28.03 ± 7.14 mW) (**Figure 4E**). Assessment of current activation kinetics with CYLE (6 μM) revealed that, whereas DOR-mediated currents activated with kinetics similar to the MOR currents previously observed in LC neurons, somato-dendritic MOR currents in CA1 PV-BCs activated 3-fold more slowly, similar to the rate observed for presynaptic MOR in these neurons (ACSF: 275.9 ± 35.7 ms, n = 11 cells; CTOP: 395.3 ± 109.6 ms, n = 6 cells; TIPP-Psi: 844.1 ± 105.2 ms, n = 9 cells; p = 0.0003, Ordinary one-way ANOVA with Tukey’s multiple comparisons test) (**Figure 4F, G**) (Ingram *et al.*, 1997; Banghart and Sabatini, 2012). The small MOR-mediated currents, coupled with similarly slow signaling kinetics in both the presynaptic and somato-dendritic compartments, suggest that MOR signaling is relatively inefficient in CA1 PV-BCs.

**Figure 4.**
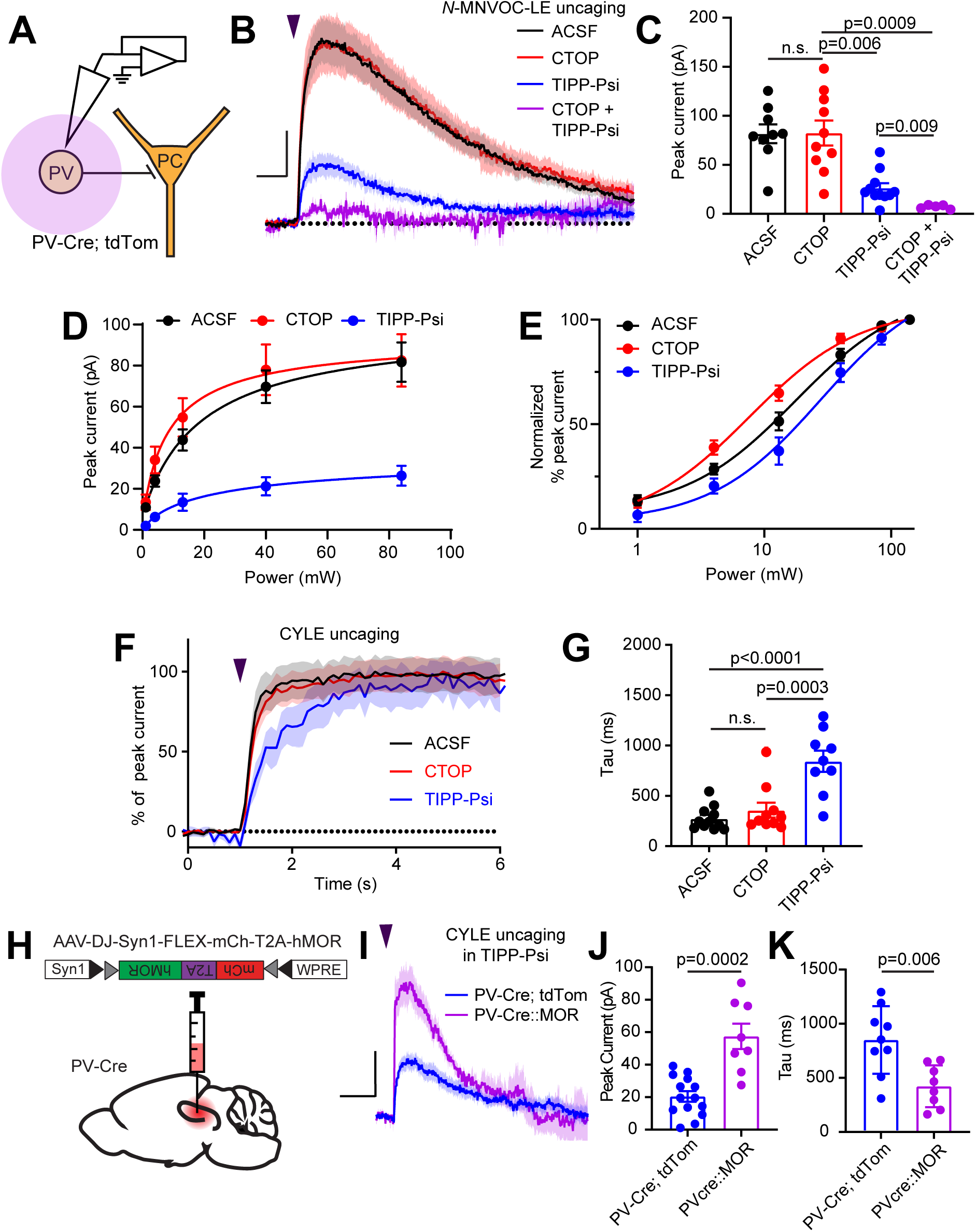
Enkephalin evokes outward currents in CA1 parvalbumin interneurons through both mu and delta opioid receptors. **A.** Schematic of whole-cell voltage clamp recording configuration from PV interneurons with peptide uncaging. **B.** Average outward currents evoked by photoactivation of *N*-MNVOC-LE (6μM) with an 84 mW light flash in the absence (black, ACSF, n = 9 cells from 5 mice) and presence of mu and delta opioid receptor antagonists (red, CTOP, n = 10 cells from 6 mice; blue, TIPP-Psi, n = 11 cells from 6 mice; purple, CTOP + TIPP-Psi, n = 5 cells from 3 mice). Scale bar: x = 5 sec, y = 20 pA. **C.** Summary of peak current amplitudes shown in **B**. **D.** Linear optical dose-response curve of peak current as a function of light intensity, in the absence (ACSF, black, n = 9 cells per laser intensity) and presence of either CTOP (red, n = 10 cells) or Tipp-Psi (blue, n = 11 cells) **E.** Logarithmic optical dose-response curves of the data in **D** normalized to the maximal peak current observed in each condition. **F.** Rising phase of the average peak-normalized outward currents evoked by photoactivation of CYLE (6μM) with an 84mW light flash in the absence (black, ACSF, n = 11 cells from 4 mice) and presence of mu and delta opioid receptor antagonists (red, CTOP, n = 10 cells from 4 mice; blue, TIPP-Psi, n = 12 cells from 4 mice). **G.** Time constants of current activation in response to photoactivation of CYLE from **F**. **H.** Schematic of viral Cre-dependent mu opioid receptor over-expression in CA1 of PV-Cre mice. **I.** Average outward currents evoked by photoactivation of CYLE by an 84 mW light flash in the presence of TIPP-Psi in either PV-Cre; tdTom mice (blue, data from **B**) or PV-Cre mice overexpressing the mu opioid receptor (purple, n = 8 cells from 3 mice). Scale bar: x = 10 sec, y = 20 pA. **J.** Summary of current amplitudes shown in **I**. **K.** Time constants of current activation in response to photoactivation of CYLE.

To identify the ion channels underlying the MOR- and DOR-mediated outward currents, we applied the GIRK channel blocker Ba^2+^ (1 mM) while delivering strong light flashes to uncage *N*-MNVOC-LE, in the absence and presence of CTOP or TIPP-Psi. Consistent with a primary role of GIRK channels, Ba^2+^ blocked the majority, but notably not all, of the current mediated by both MOR and DOR (**Supporting Figure S2A, B**). Inclusion of the HCN channel blocker ZD7288 (1 μM) further inhibited the DOR-current, but did not abolish it, suggesting the involvement of additional ion channels (Ba^2+^ in ACSF: 67.9 ± 4.9 %, n = 8 cells; Ba^2+^ in CTOP: 59.6 ± 9.7 %, n = 10 cells; Ba^2+^, Zd7288 in CTOP: 74.0 ± 5.6 %, n = 9 cells; Ba^2+^ in TIPP-Psi: 67.7 ± 9.1 %, n = 11 cells; no significant differences, Ordinary one-way ANOVA).

One possible explanation for the slow kinetics and low efficacy of MOR-mediated GIRK activation, as well as slow kinetics of synaptic suppression, is relatively low cell surface expression of MORs in comparison to DORs. In LC, reducing available surface MORs with a covalent antagonist leads to a reduction not only in the amplitude of MOR-mediated currents, but also a slowing of activation kinetics (Williams, 2014). To test this hypothesis, we virally overexpressed human MOR (hMOR) with an mCherry tag in PV-Cre mice and probed the resulting enhanced MOR signaling with CYLE in TIPP-Psi (Liu *et al., in press*) (**Figure 4H, I**). As predicted, hMOR overexpression enhanced both the magnitude (57.5 ± 7.8 pA, n = 8 cells, p = 0.0002, Mann-Whitney test) and the kinetics (421.8 ± 68.7 ms, n = 8 cells, p = 0.006, Mann-Whitney test) of the MOR-mediated current evoked with a strong light flash in comparison to those recorded from PV-Cre; tdTom mice (**Figure 4I-K**). Both parameters correlated strongly with mCherry fluorescence as an indicator of expression level (Peak: r = 0.8314, Tau on: r = −0.8538, Pearson’s correlation coefficient) (**Supporting Figure S2C, D**). These results indicate that low MOR expression levels can account for the surprisingly modest effects of MOR activation in the somato-dendritic compartment of PV-BCs.

### MORs and DORs do not functionally interact in CA1 PV-BCs

The apparent co-expression of MORs and DORs in the somato-dendritic compartment is a minimal requirement for functional interactions between receptors. We therefore asked if MORs and DORs undergo heterologous desensitization such that desensitization of one receptor perturbs the function of the other. We first confirmed that prolonged exposure to DAMGO (1 μM) caused desensitization of the resulting outward current (**Figure 5A**). After incubating slices in DAMGO for at least 10 minutes to maximally desensitize MOR, dose-response curves were obtained in the presence of DAMGO, such that subsequent photorelease of LE would only activate DORs (**Figure 5B**). We compared these responses to those evoked in naïve slices bathed in the MOR antagonist CTOP. Indicative of a lack of heterologous desensitization, neither the efficacy or potency of LE at DORs was affected by MOR desensitization (EC50 value of LE in the presence of DAMGO: 5.12 ± 0.38 mW, n = 9 cells; CTOP: 6.10 ± 0.53 mW, n = 7 cells) (**Figure 5C, D**). Similarly, prolonged exposure to deltorphin II (1 μM) caused desensitization of the outward current (**Figure 5E**). Desensitization of DORs using deltorphin II did not affect the ability of LE to elicit somato-dendritic outward currents compared to naïve slices bathed in the DOR antagonist TIPP-Psi (EC50 value of LE in the presence of Delt II: 13.47 ± 1.11 mW, n = 7 cells; TIPP-Psi: 12.55 ± 1.8 mW, n = 9 cells). These results reveal that MORs and DORs do not undergo heterologous desensitization in CA1 PV-BCs.

**Figure 5.**
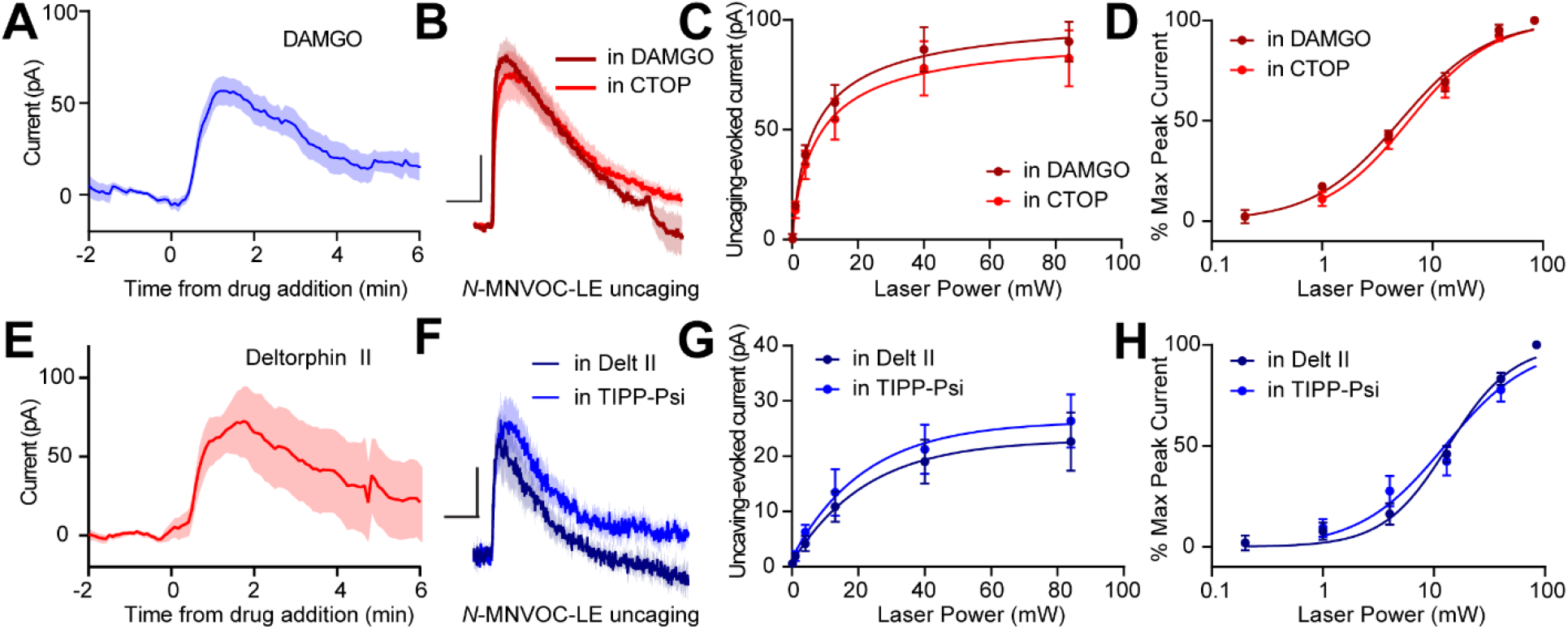
Somato-dendritic mu and delta opioid receptors do not exhibit heterologous desensitization. **A.** Average outward current evoked by sustained bath application of DAMGO (n = 9 cells from 6 mice). **B.** Average outward currents evoked by photoactivation of *N*-MNVOC-LE either in the presence of CTOP (red, data from **4B**) or in the presence of DAMGO, after desensitization (brick red, n = 9 cells from 4 mice). Scale bars: x = 10 sec, y = 25 pA. **C.** Linear optical dose-response curve of peak current as a function of light intensity, in the presence of either CTOP (red, n = 10 cells, data from **4C**) or DAMGO (brick red, n = 9 cells) **D.** Logarithmic optical dose-response curves of the data in **C** normalized to the maximal peak current observed in each condition. **E.** Average outward current evoked by sustained bath application of deltorphin II (n = 12 cells from 6 mice). **F.** Average outward currents evoked by photoactivation of *N*-MNVOC-LE either in the presence of TIPP-Psi (blue, data from **4B**) or in the presence of deltorphin II, after desensitization (navy blue, n = 8 cells from 4 mice). Scale bars: x = 10 sec, y = 10 pA. **G.** Linear optical dose-response curve of peak current as a function of light intensity, in the presence of either TIPP-Psi (blue, n=11 cells, data from **4C**) or deltorphin II (navy blue, n = 8 cells) **H.** Logarithmic optical dose-response curves of the data in **F** (blue, TIPP-Psi, n = 9 cells; navy blue, deltorphin II, n = 7 cells) normalized to the maximal peak current observed in each condition.

MORs and DORs have been proposed to functionally interact through the formation of heteromeric receptors such that a selective antagonist for one receptor enhances signaling at the other (Gomes *et al.*, 2004). To directly probe for functional interactions of this type, we developed a new photoactivatable analogue of the MOR-selective agonist DAMGO, CNV-Y-DAMGO (Ma et al, *in preparation*). We hypothesized that if these interactions are present, inclusion of TIPP-Psi in the bath would lead to a leftward shift in the optical dose-response curves of CNV-Y-DAMGO, and possibly an increase in the response kinetics. We tested this by uncaging CNV-Y-DAMGO (1 μM) while measuring somato-dendritic currents in PV-BCs (**Figure 6A-E**) and eIPSCs in pyramidal neurons (**Figure 6F-J**). In both cases, TIPP-Psi did not alter either the kinetics of the response to DAMGO photorelease (GIRK tau on CNV-Y-DAMGO: 917.6 ± 75.7 ms, n = 11 cells; CNV-Y-DAMGO + TIPP-Psi: 808.8 ± 46.5 ms, n = 7 cells; no significant difference, t-test; eIPSC tau on CNV-Y-DAMGO: 476.4 ± 36.9 ms, n = 8 cells; CNV-Y-DAMGO + TIPP-Psi: 441.6 ± 28.1 ms, n = 7 cells; no significant difference, t-test) (**Figure 6C, H**), its maximal effect (**Figure 6D, I**), or its dose-dependence (EC50 values for GIRKs in CNV-Y-DAMGO: 6.86 ± 0.68 mW, n = 8 cells; CNV-Y-DAMGO + TIPP-Psi: 8.53 ± 0.64 mW, n = 7 cells; EC50 values for eIPSCs in CNV-Y-DAMGO: 2.79 ± 0.44 mW, n = 9 cells; CNV-Y-DAMGO + TIPP-Psi: 3.06 ± 0.38 mW, n = 9 cells)(**Figure 6E, J**). These results indicate that MORs and DORs do not interact in PV-BCs in a manner consistent with MOR/DOR heteromers.

**Figure 6.**
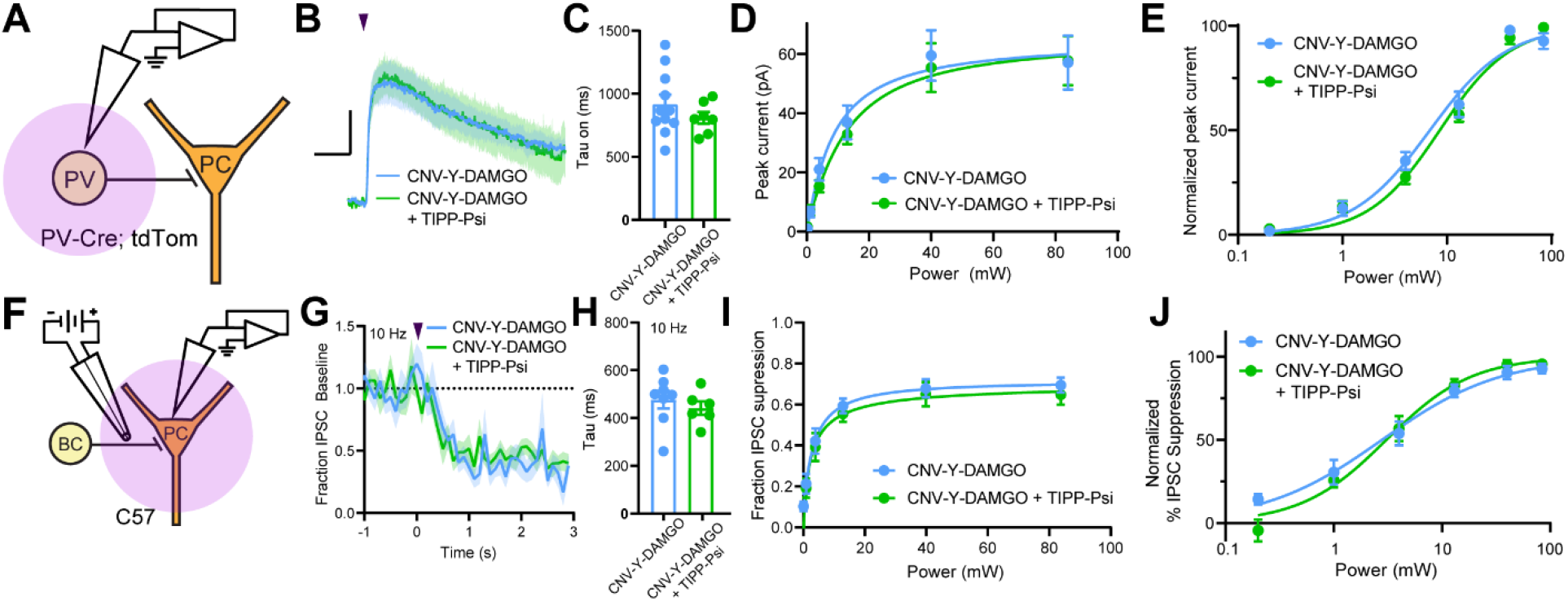
Mu and delta opioid receptors do not signal as heterodimers in CA1 PV neurons. **A.** Schematic of whole-cell voltage clamp recording configuration from PV interneurons with peptide uncaging. **B.** Average outward currents evoked by photoactivation of CNV-Y-DAMGO with an 84 mW light flash either in the absence (sky blue, n = 8 from 5 mice) or presence (green, n = 7 cells from 4 mice) of TIPP-Psi. Scale bar: x = 10 sec, y = 20 pA. **C.** Time constants of current activation in response to photoactivation of CNV-Y-DAMGO in the absence or presence of TIPP-Psi. **D.** Linear optical dose-response curve of peak current as a function of light intensity, in the absence (sky blue) or presence (green) of TIPP-Psi. **E.** Logarithmic optical dose-response curves of the data in **D** normalized to the maximal peak current observed in each condition. **F.** Schematic of the experimental configuration for photo-uncaging of opioid neuropeptides while recording electrically-evoked inhibitory synaptic transmission in wild type mice. **G.** Average, baseline subtracted and baseline-normalized IPSC amplitude showing the kinetics of synaptic suppression with electrical stimulation at 10 Hz in the absence (sky blue, n = 8 cells from 4 mice) or presence of TIPP-Psi (green, n = 8 cells from 4 mice). **H.** Time constants of synaptic suppression at 10 Hz stimulation in response to photoactivation of CNV-Y-DAMGO in the absence or presence of TIPP-Psi. **I.** Linear optical dose-response curve of IPSC suppression as a function of light intensity, in the absence (sky blue) or presence (green) of TIPP-Psi. **J.** Logarithmic optical dose-response curves of the data in **I** normalized to the maximal IPSC suppression observed in each condition.

## Discussion

### Identification of the delta opioid receptor as the primary target of enkephalin in CA1 PV-BCs

Prior models of neuromodulator actions on hippocampal interneurons have emphasized MOR expression as a primary distinctive feature of PV-BCs, as opposed to CCK-BCs (Freund and Katona, 2007). This results from an electrophysiological study in CA1 BCs that used the MOR agonist DAMGO to elicit outward somato-dendritic currents and suppress synaptic output (Glickfeld, Atallah and Scanziani, 2008). Although multiple studies have demonstrated the expression of DORs, in addition to MORs, in CA1 PV neurons, the relative contributions of the two receptors to opioid modulation of CA1 PV-BCs has not been established (Stumm *et al.*, 2004; Erbs *et al.*, 2012; Faget *et al.*, 2012). Our findings, using enkephalin to activate both MORs and DORs, indicate that DORs dominate cellular and synaptic responses to enkephalin, in particular at low concentrations that may be most physiologically relevant. Optically-generated doseresponse curves with caged enkephalin revealed that LE activates DORs with ~3-fold greater potency than MORs in both the somato-dendritic and presynaptic compartments. Strikingly, the dose-response relationships observed in the absence of antagonist closely match those obtained with MORs blocked, which underscores the dominant role of DORs in the integrated response to enkephalin. While this may reflect a greater binding affinity of LE for DORs (Toll *et al.*, 1998), because somato-dendritic DOR-mediated currents are much larger than MOR-mediated currents when both receptors are saturated, this preferential recruitment of DOR signaling is also likely to result in much stronger inhibition of cellular excitability. In presynaptic terminals of PV-BCs, the strong reciprocal occlusion of synaptic suppression by saturating doses of selective MOR and DOR agonists suggests that because DOR activation by LE occurs at lower concentrations, it will occlude subsequent actions of MOR at higher doses. Given that local sources of the MOR-selective neuropeptide β-endorphin are apparently lacking in CA1 (Bjorklund and Hokfelt, 1986), this raises the question as to why PV-BCs express MORs at all. One possible explanation is that diurnal variation in the levels of brain-wide β-endorphin in the cerebrospinal fluid contribute to the resting excitability and tune the strength of synaptic output via PV-BC MORs, while dynamic, local release of enkephalin in CA1 produces stronger, temporally-precise inhibition of cellular output through activation of DORs (Dent *et al.*, 1981; Barreca *et al.*, 1986). Resolving this apparent conflict will require the identification of behavior contexts that result in endogenous enkephalin release in CA1, as well as the sources that provide enkephalin to presynaptic and somato-dendritic opioid receptors in PV-BCs.

### Enkephalin suppresses synaptic transmission with sub-second kinetics

Although GPCRs are well established to engage effector pathways within 100 ms of exposure to agonists, data describing the kinetics of synaptic suppression by Gα_i/o_-coupled GPCRs are sparse. A study in rat cerebellum reported rapid and transient GABAB-mediated suppression of an excitatory synapse peaked 300 ms after application of a high frequency stimulus to drive GABA release, with detectable reduction in presynaptic Ca^2+^ 100 ms after the stimulus (Dittman and Regehr, 1997). A similarly structured study in rat striatum observed a maximal suppression of corticostriatal transmission 500 ms after stimulating striatal neurons to release endogenous opioid neuropeptides (Blomeley and Bracci, 2011). Both of these studies involved relatively small quantities of neuromodulator such that rapid clearance likely obscured the intrinsic kinetics of the presynaptic signaling pathway. Here, we found that photorelease of enkephalin during high frequency stimulation of synaptic transmission produced suppression that peaked between 1-2 s after the light flash. The high sample frequency we employed facilitated rate determination, yielding an average time constant of ~300 ms at 10 Hz. A potential caveat to our approach is that our measurements were taken from synapses that were already in a partially depressed state. Nonetheless, we observed a striking difference in the kinetics of synaptic suppression by DORs and MORs that closely matched the time constants determined for the activation of outward current in the somato-dendritic compartment. In both cases, MORs exhibited much slower kinetics (tau ~800 ms) than DORs. This was not ligand-dependent, as the same time constants were obtained using caged DAMGO (**Figure 6C, H**). This stands in contrast to prior measurements of the kinetics of GIRK activation by MORs in other cell types that found faster time constants, similar to our measurements of DOR-mediated responses (Ingram *et al.*, 1997; Banghart and Sabatini, 2012; Williams, 2014). Interestingly, in the somato-dendritic compartment, we found that increasing MOR expression increased the MOR-evoked current activation rate. Thus, differences in MOR kinetics observed for other brain regions or cell types is likely to reflect differences in relative levels of MOR expression.

It is also notable that relatively strong activity-dependent synaptic depression due to high frequency stimulation did not dramatically occlude synaptic suppression, indicating that release of a relatively depleted readily-releasable pool of vesicles is still prone to attenuation by Gα_i/o_-coupled GPCRs that inhibit presynaptic Ca2^+^ channels. We observed a modest but significant negative correlation between the extent of synaptic suppression and the frequency of stimulation, which is consistent with voltage-dependent unbinding of Gβγ from VSCCs (Bean, 1989; Brody *et al.*, 1997).

### Lack of functional interactions between MORs and DORs in CA1 PV-BCs

MORs and DORs have been suggested to physically interact via the formation of heterodimers when expressed in the same cell. Although most of the mechanistic work on MOR/DOR heteromers has been performed in cultured cells with overexpressed receptors, multiple studies have also found evidence for their occurrence in naïve brain tissue (Gomes *et al.*, 2004; Gupta *et al.*, 2010; Kabli *et al.*, 2014; Erbs *et al.*, 2015). The pharmacological framework for detecting MOR/DOR functional interactions emerges from studies in cultured cells showing that ligands for one receptor can increase the binding (in terms of B_max_ but not K_d_) and signaling efficacy of agonists for the other (Gomes *et al.*, 2000). Specifically, both the DOR selective agonist deltorphin II and the selective antagonist TIPP-Psi were observed to enhance binding of DAMGO, which was accompanied by a decrease in DAMGO’s EC_50_ in a functional assay of MOR activation. Conversely, DAMGO, as well as the MOR antagonist CTOP, enhanced binding and reduced the EC_50_ of deltorphin II. Similar enhancements of MOR activation in the presence of DOR antagonist have been observed in brain tissue using multiple functional assays of MOR signaling, including antinociceptive behavior (Gomes *et al.*, 2004).

Additional evidence supporting the existence of endogenous MOR/DOR heteromers has emerged from the observation that the efficacy of bivalent MOR-DOR ligands is highly dependent on the length of the linker connecting them, which is consistent with action at a receptor complex (Daniels *et al.*, 2005). Numerous studies of receptor trafficking in cultured cells indicate substantial co-localization of MORs and DORs, as well as co-internalization upon exposure to certain agonists for one of the two receptors (*e.g.* He *et al.*, 2011; Derouiche *et al.*, 2020). In addition, biochemical studies have reported co-immunoprecipitation from naïve brain tissue using an antibody for either MORs or DORs (Gomes *et al.*, 2000), or an antibody that specifically recognizes MOR/DOR heteromers (Gupta *et al.*, 2010).

In contrast to these prior studies, we found no evidence for functional interactions between MORs and DORs in CA1 PV-BCs. Rather than synergistic, supralinear signaling, we observed largely parallel signaling and occlusion. If LE elicited synergistic signaling between MORs and DORs, we would predict that the dose-response curve for LE with both receptors intact (control conditions) would sit to the left of the curves obtained for either receptor in isolation using selective antagonists. This was not the case. Instead, in both subcellular compartments, DOR activation accounted for the low end of the dose-response curves, with MORs contributing only at higher concentrations. Strong occlusion at presynaptic terminals was observed, as simultaneous application of small molecule agonists for both receptors only slightly increased the extent of synaptic modulation in comparison to either drug alone (from 70% to ~75% suppression). Similar occlusion was also observed while monitoring presynaptic Ca^2+^ transients. Interestingly, only unidirectional occlusion was observed in the somato-dendritic compartment, where MOR block had no effect on outward currents driven by high doses of LE, while DOR block dramatically reduced them. This observed sub-linear signaling indicates that DORs have access to a larger pool of GIRKs than MORs, and that GIRKs activated by MORs are completely shared between both receptor types. These results are summarized graphically in **Figure 7**.

**Figure 7.**
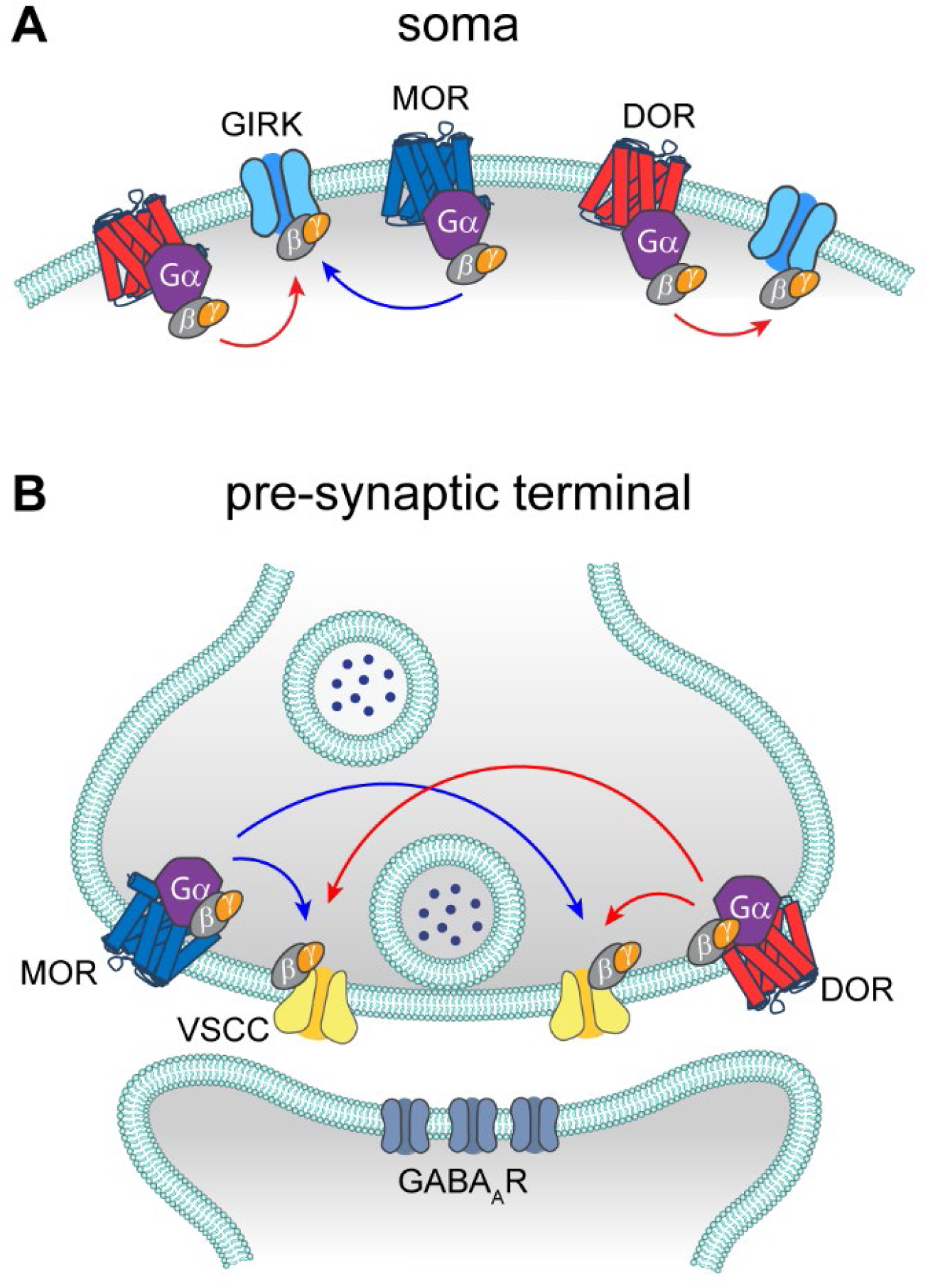
Models of MOR and DOR signaling in the soma and the pre-synaptic terminal. **A.** In the soma, both MORs (blue) and DORs (red) signal through GIRK channels. MORs are expressed at lower levels than DORs, as the somato-dendritic currents evoked by activation of MORs alone are small and are increased by increasing MOR expression. The unidirectional occlusion observed suggests that MORs only have access to a subset of GIRKs, whereas DORs have access to larger pool that encompasses the MOR-pool. **B.** In the pre-synaptic terminal, MORs and DORs both act on VSCCs to suppress Ca^2+^ influx and inhibit vesicle release. Unlike somatic MORs and DORs, pre-synaptic MORs and DORs are bidirectionally occlusive, so that both MORs and DORs have access to the majority of VSCCs.

In addition, we did not observe heterologous desensitization between MORs and DORs. In opioid-naïve animals, desensitization appears to occur at the level of the receptor, likely due to C-terminus phosphorylation, rather than through the effectors (Llorente *et al.*, 2012; Leff, Arttamangkul and Williams, 2020). Nonetheless, because desensitization can lead to endocytosis, and possibly conformational changes, if the receptors were physically interacting, desensitization of one receptor may be expected to impact signaling at the other.

Similarly, our findings argue against the presence of native MOR/DOR heteromers in either the somato-dendritic or presynaptic compartments of CA1 PV-BCs, since TIPP-Psi had no effect on DAMGO potency or signaling kinetics, both of which serve as sensitive measures of receptor function. This lack of interaction between MORs and DORs is consistent with our previous observation in striatal indirect pathway neurons, wherein their actions were strictly additive, and genetic removal of either receptor neither enhanced nor suppressed the efficacy of the other (Banghart *et al.*, 2015). A possible explanation is that MOR/DOR heteromers present in PV-BCs are retained in the Golgi apparatus due to a lack of Rtp4 expression (*Allen Brain Atlas API;* Décaillot *et al.*, 2008; Saunders *et al.*, 2018). As this may involve sequestering MORs, it may also contribute to the surprisingly small somato-dendritic MOR-mediated GIRK currents we observed. While MOR/DOR functional interactions may be more prominent in other brain regions, our findings indicate that co-expression and co-localization in subcellular compartments do not guarantee receptor crosstalk at the cell surface.

In conclusion, DORs in CA1 PV-BCs, rather than MORs, are the primary target of the opioid neuropeptide enkephalin. Although signaling at both receptors converges on largely overlapping populations of effectors within the same subcellular compartments, MORs and DORs appear to signal predominantly in a parallel, functionally-independent manner. These results imply that functional redundancy between multiple GPCRs expressed in the same neuron may be a common feature in the nervous system.

Additional research is necessary to further delineate mechanisms that determine whether or not heteromers form when heterophilic receptors are present in close proximity within cells.

## Materials and Methods

### Key Resources Table

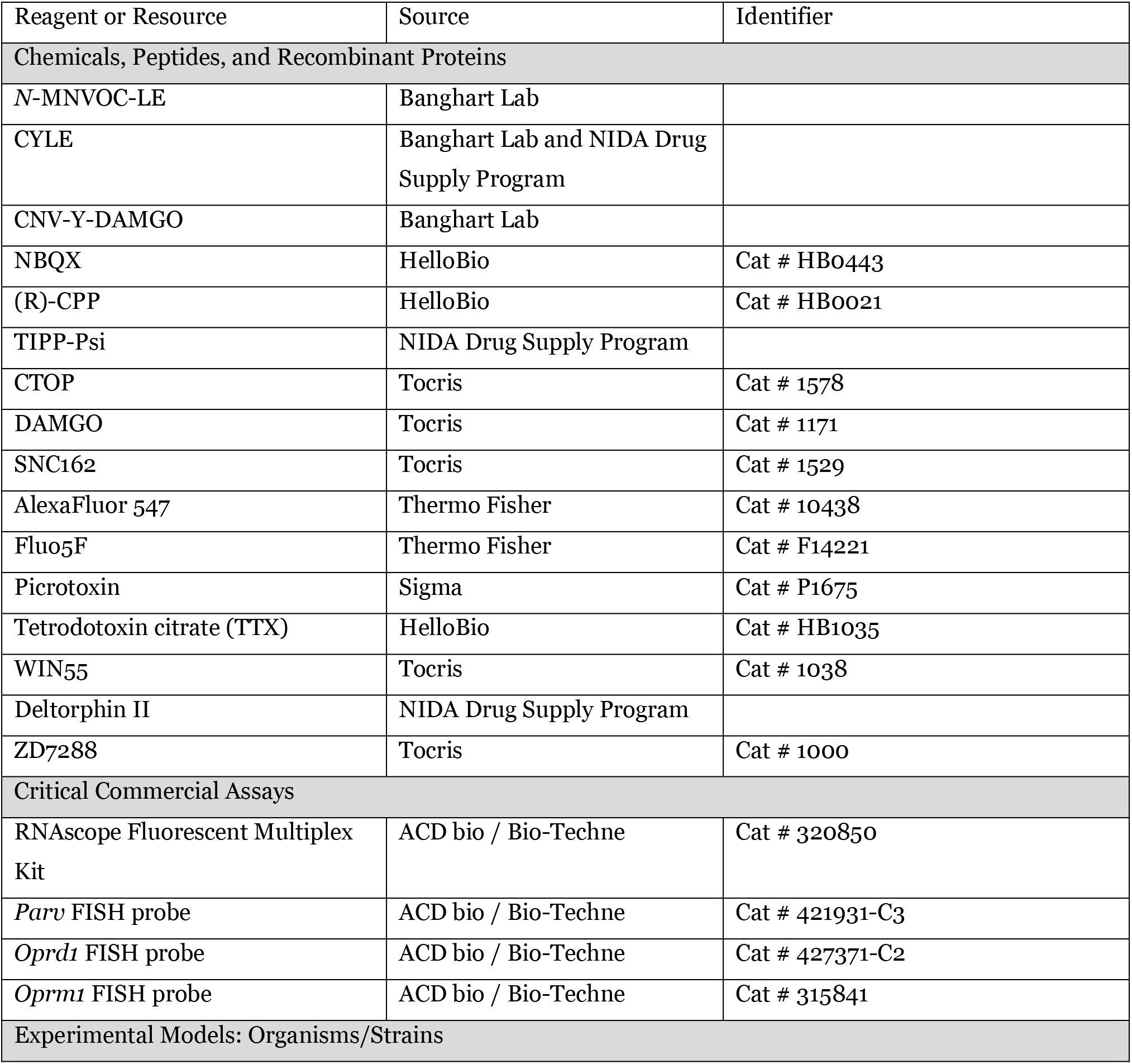
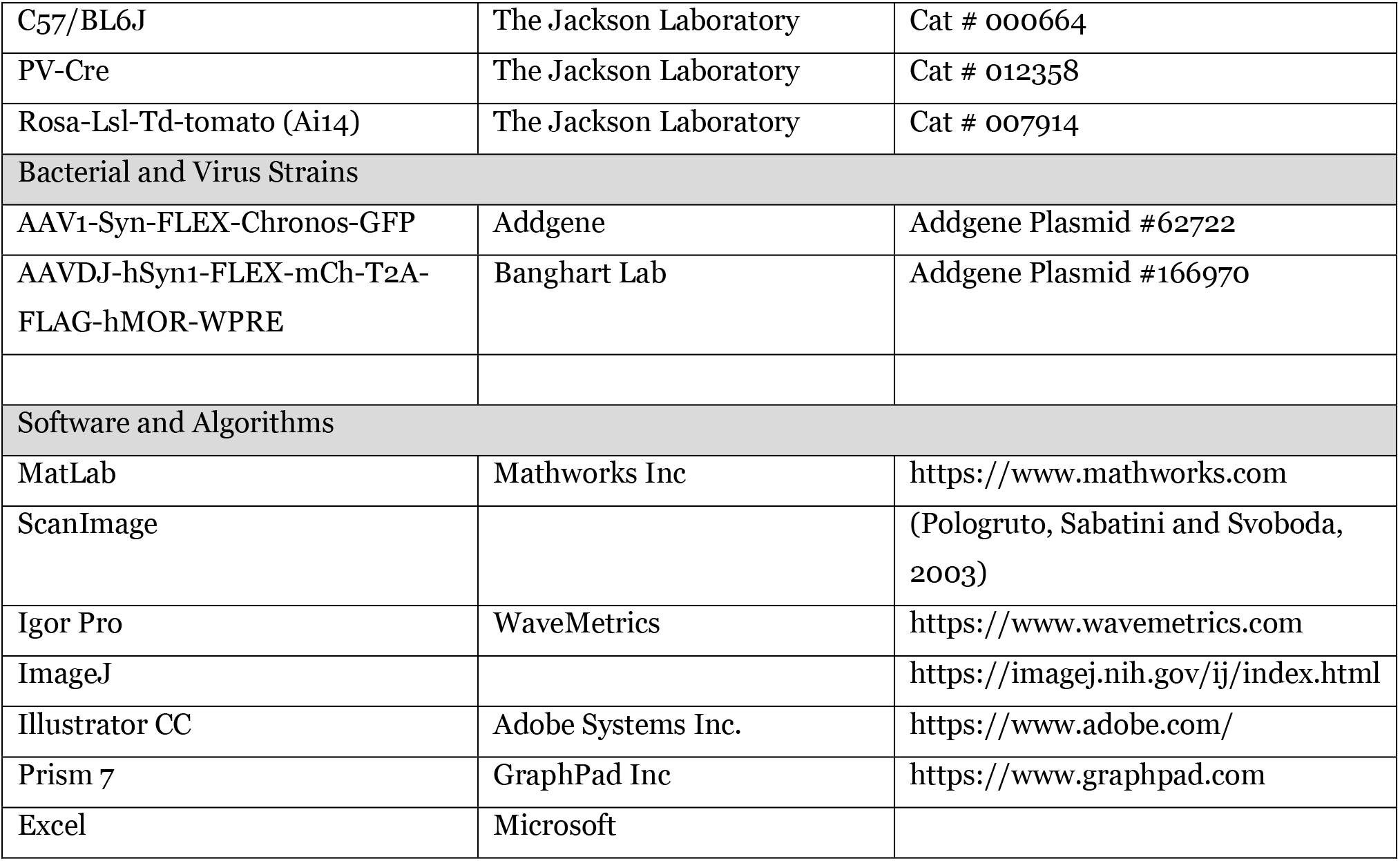

### Brain Slice Preparation

Animal handling protocols were approved by the UC San Diego Institutional Animal Care and Use Committee. Postnatal day 15-35 mice on a C57/Bl6 background were anesthetized with isoflurane and decapitated, and the brain was removed, blocked, and mounted in a VT1000S vibratome (Leica Instruments). Horizontal slices (300 μm) were prepared in ice-cold choline-ACSF containing (in mM) 25 NaHCO3, 1.25 NaH2PO4, 2.5 KCl, 7 MgCl2, 25 glucose, 0.5 CaCl2, 110 choline chloride, 11.6 ascorbic acid, and 3.1 pyruvic acid, equilibrated with 95% O2/5% CO_2_. Slices were transferred to a holding chamber containing oxygenated artificial cerebrospinal fluid (ACSF) containing (in mM) 127 NaCl, 2.5 KCl, 25 NaHCO3, 1.25 NaH2PO4, 2 CaCl2, 1 MgCl2, and 10 glucose, osmolarity 290. Slices were incubated at 32 °C for 30 min and then left at room temperature until recordings were performed.

### Electrophysiology

All recordings were performed within 5 h of slice cutting in a submerged slice chamber perfused with ACSF warmed to 32 °C and equilibrated with 95% O_2_/5% CO_2_. Whole-cell voltage clamp recordings were made with an Axopatch 700B amplifier (Axon Instruments). Data were filtered at 3 kHz, sampled at 10 kHz, and acquired using National Instruments acquisition boards and a custom version of ScanImage written in MATLAB (Mathworks). Cells were rejected if holding currents exceeded −200 pA or if the series resistance (<25 MΩ) changed during the experiment by more than 20%. For recordings measuring K+ currents in PV cells (Figure 1), patch pipets (open pipet resistance 2.0-3.0 MΩ) were filled with an internal solution containing (in mM) 135 KMeSO_4_, 5 KCl, 5 HEPES, 1.1 EGTA, 4 MgATP, 0.3 Na2GTP, and 10 Na2phosphocreatine (pH 7.25, 286 mOsm/kg). Cells were held at −55 mV, and synaptic transmission was blocked with the addition to the ACSF of 2,3-dihydroxy-6-nitro-7-sulfamoyl-benzo(f)quinoxaline (NBQX; 10 μM), R,S-3-(2-carboxypiperazin-4-yl)propyl-1-phosphonic acid (CPP; 10 μM), picrotoxin (10 μM), and TTX (1uM). For recordings measuring inhibitory synaptic transmission in mouse hippocampus, patch pipets (2.5-3.5 MΩ) were filled with an internal solution containing (in mM) 135 CsMeSO3, 10 HEPES, 1 EGTA, 3.3 QX-314 (Cl - salt), 4 Mg-ATP, 0.3 Na-GTP, and 8 Na 2 phosphocreatine (pH 7.3, 295 mOsm/kg). Cells were held at 0 mV to produce outward currents. Excitatory transmission was blocked by the addition to the ACSF of NBQX (10 μM) and CPP (10 μM).To electrically evoke IPSCs, stimulating electrodes pulled from theta glass with ~5 μm tip diameters were placed at the border between stratum pyramidale and straum oriens nearby the recorded cell (~50-150 μm) and a two brief pulses (0.5 ms, 50-300 μA, 50 ms interval) were delivered every 20 s.

### UV Photolysis

Uncaging was carried out using 5 ms flashes of collimated full-field illumination with a 355 nm laser, as previously described. Light powers in the text correspond to measurements of a 10 mm diameter collimated beam at the back aperture of the objective. Beam size coming out of the objective onto the sample was 3900 μm^2^.

### Optogenetics

AAV encoding Chronos-GFP was injected into the hippocampus of PV-Cre pups P0-3. The virus was allowed to express for 4 weeks and then acute hippocampal slices were made as described above. For optogenetic stimulation of PV basket cell terminals, two 2 ms pulses of blue LED were flashed over the cell body of the patched pyramidal cell. The field stop of the LED was narrowed to 6600 μm^2^ in order to limit the excitation to only the immediate axons surrounding the cell body.

### Two-photon calcium imaging

Two-photon imaging of axonal boutons was performed using a custom-built two-photon laser-scanning microscope (Carter and Sabatini, 2004; Bloodgood and Sabatini, 2007). First, PV neurons in the CA1 region of the hippocampus were visualized using epifluorescence in a PV-Cre; tdTom line and targeted recordings were made under infrared differential interference contrast (IR-DIC) on an Olympus BX51 microscope. Whole cell current clamp recordings were made with a potassium (K)-methanesulfonate internal consisting of (in mM): 135 KMeSO_4_, 5 KCl, 5 HEPES, 4 MgATP, 0.3 Na2GTP, and 10 Na2phosphocreatine. The internal also contained the Ca-sensitive green fluorophore Fluo-5F (300μM) and Ca-insensitive red fluorophore Alexa Fluor-594 (30μM). After a patch was made, the cell was allowed at least 15 minutes for the dye and indicator to fill the axons. Then an 800nm laser was used to locate axonal boutons based on morphology. Once identified, line scans were made across 1-2 boutons while evoking 1 or 5 action potentials by injecting voltage into the cell body. Calcium transients were averaged across 30 trials, before and after drug addition. Stimulus-evoked changes in fluorescence (and the Ca signal) were reported as %ΔG/Gsat, reflecting measurements of ΔG/R normalized to G/R in saturating Ca as described previously (Bloodgood and Sabatini, 2007).

### Data Analysis

Electrophysiology data were analyzed in Igor Pro (Wavemetrics). Peak current amplitudes were calculated by averaging over a 200 ms (GIRK) or 2 ms (synaptic transmission) window around the peak. Activation time constants for GIRKs were calculated by fitting the rising phases of light evoked currents to an exponential function. To determine magnitude of modulation by enkephalin uncaging (%IPSC suppression), the IPSC peak amplitude immediately after a flash was divided by the average peak amplitude of the three IPSCs preceding the light flash. Kinetics of synaptic modulation (Figure 3) were determined by averaging 3 stimulus trains before uncaging (at 10 Hz, 20 Hz, and 50 Hz) and fitting a bi-exponential curve describe the synaptic depression. The curve was then divided from the stimulus train with uncaging to get the traces seen in Figure 3B. The time constant was then extracted from a mono-exponential was fit to the suppression from the time of uncaging. The effects of drugs on IPSC suppression were calculated as the average %IPSC suppression 2-3 minutes after drug addition. Summary values are reported as mean ± SEM. All data are treated as parametric and specific statistical tests and corrections are described for each figure in the text and figure legends.

### Fluorescence in-situ hybridization

Mice were deeply anesthetized with isoflurane and decapitated, and their brains were quickly removed and frozen in tissue freezing medium on dry ice. Brains were cut on a cryostat (Leica CM 1950) into 8μm sections, adhered to SuperFrost Plus slides (VWR), and stored at −80°C. Samples were fixed 4% paraformaldehyde, processed according to ACD RNAscope Fluorescent Multiplex Assay manual, and coverslipped with ProLong antifade reagent (Molecular Probes). Sections were imaged on a Keyence BZ-X710 Microscope at 60x magnification. The images were acquired and manually scored for the presence of fluorescent puncta and colocalization using ImageJ.

## Acknowledgements

We thank the National Institute on Drug Abuse Drug Supply Program (NDSP) for generously providing pharmacological reagents; L Sancho and E Campbell for training and assistance with two-photon microscopy; BK Lim for reagents for adenoassociated virus production; E Berg for genotyping, animal husbandry, adenoassociated virus production and administrative assistance; J Isaacson, W Birdsong, J Williams, M Lovett-Barron, and members of the Banghart Lab for helpful discussions.

## Competing interests

The authors declare no competing interests.

## Funding

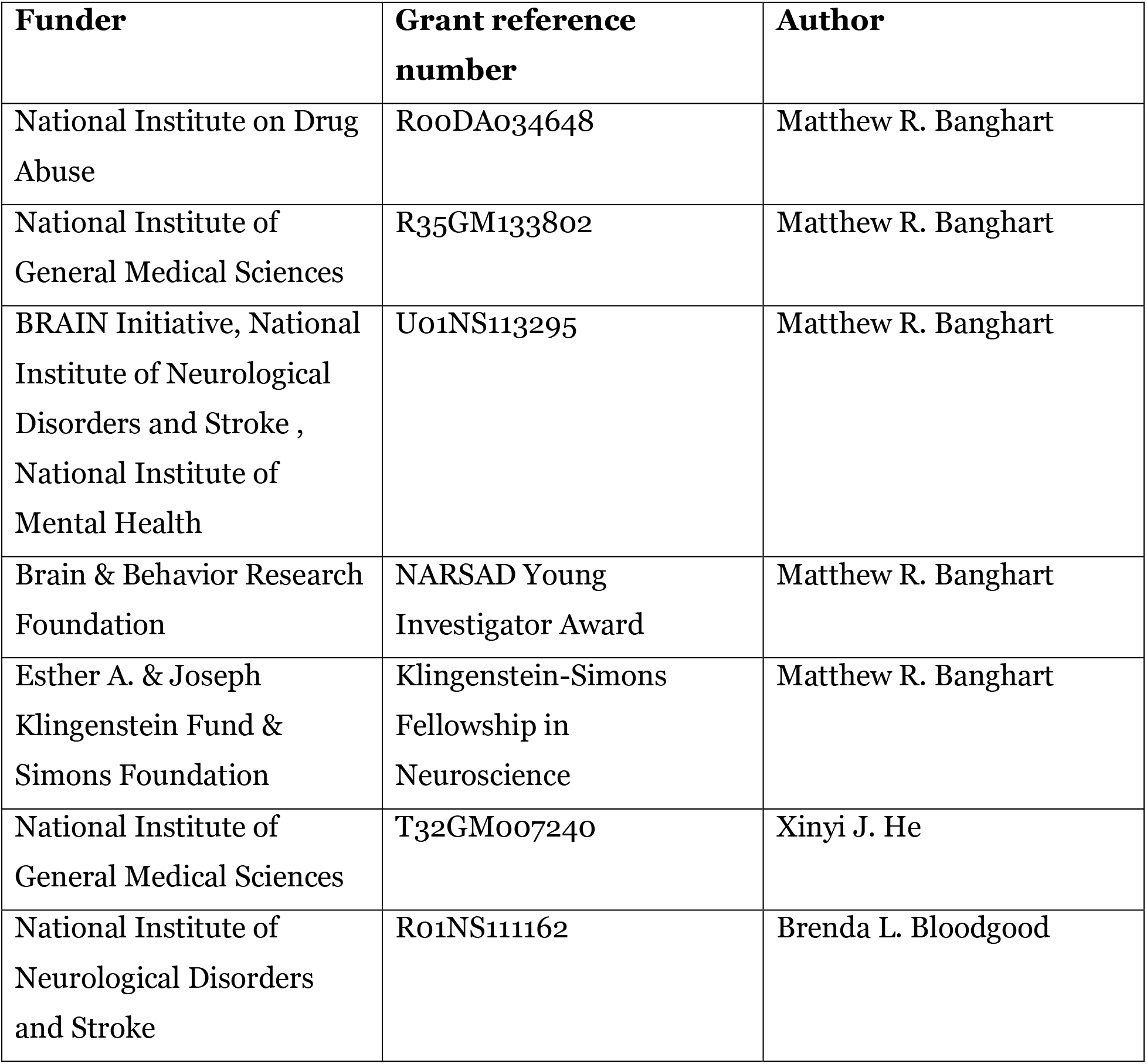

The funders had no role in study design, data collection and interpretation, or the decision to submit the work for publication

## Author contributions

Xinyi Jenny He: conceptualization, data curation, formal analysis, investigation, methodology, writing-original draft, writing-review and editing

Janki Patel, data curation, formal analysis

Connor E. Weiss, data curation

Xiang Ma, resources

Brenda L. Bloodgood, resources, methodology, writing-review and editing

Matthew R. Banghart, conceptualization, resources, funding acquisition, methodology, writing-original draft, project administration, writing-review and editing

## Ethics

Animal experimentation: All procedures were performed in accordance with protocols approved by the University of California San Diego Institutional Animal Care and Use Committee (IACUC) following guidelines described in the the US National Institutes of Health Guide for Care and Use of Laboratory Animals (UCSD IACUC protocol S16171). All surgery was performed under isoflurane anesthesia.

---

**Supporting Figure S1 (refers to Figure 1).**
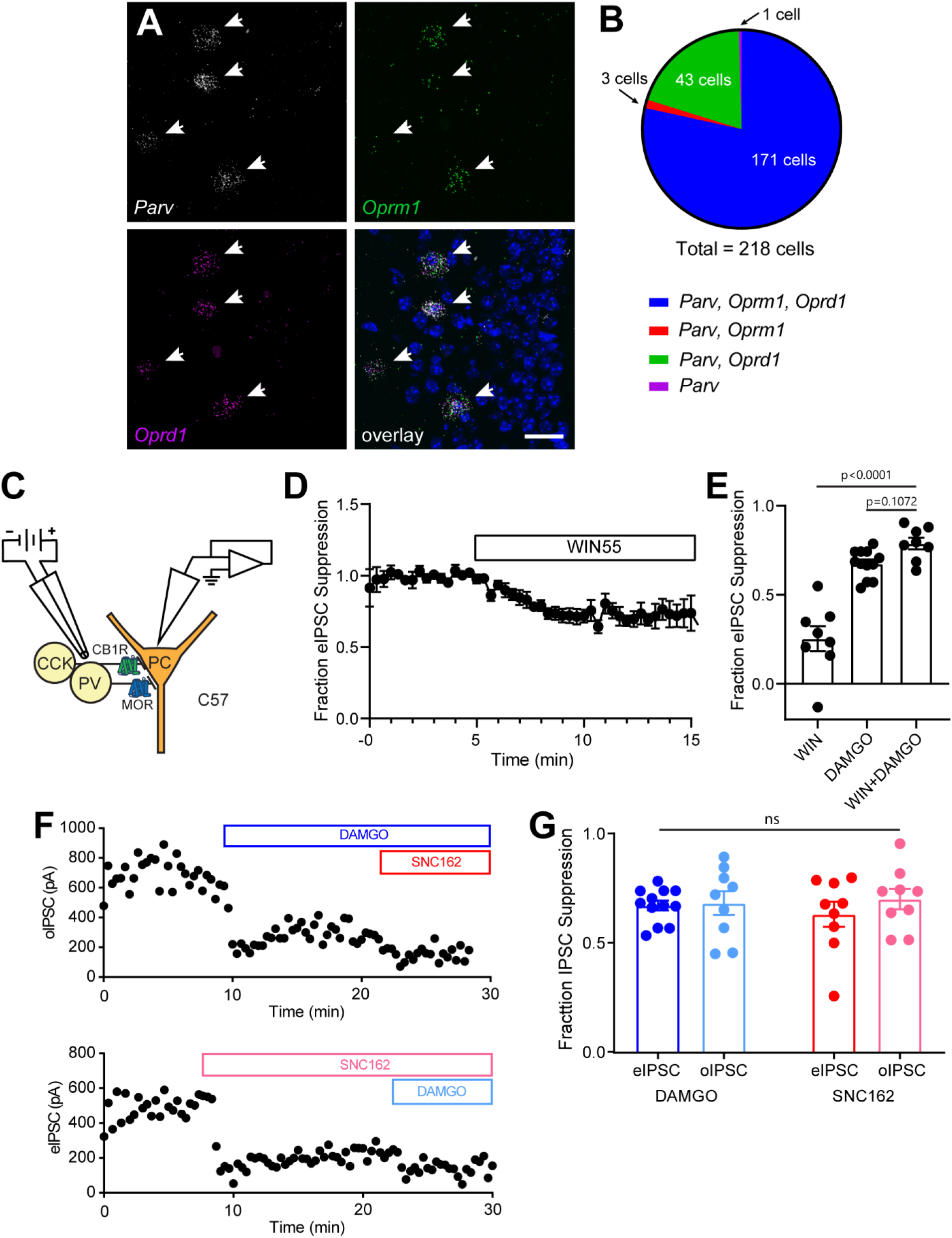
Opioid receptor mRNA in CA1 parvalbumin interneurons and characterization of the neuromodulator-sensitivity of CA1 basket cell synaptic output. **A.** Example fluorescence *in situ* hybridization image of *Parv, Oprm1,* and *Oprd1* mRNA in the CA1 pyramidal layer of mouse hippocampus. Scale bar = 20 μm. **B.** Summary of *Parv, Oprm1*, and *Oprd1* mRNA co-localization images acquired from 5 mice. **C.** Schematic of recording configuration for electrical stimulation depicting two populations of basket cells and their distinguishing neuromodulator receptors. **D.** Baseline-normalized, average eIPSC amplitude over time during bath application of the CB1R agonist WIN55 (n=9 cells from 3 mice). **E.** Summary data of WIN55 and DAMGO flow-in experiments (DAMGO data replotted from Figure 1I), revealing only a small contribution to the eIPSC from CCK-BCs that are suppressed by CB1R but not MOR. **F.** Example double flow-in experiments with optogenetic stimulation (top) and electrical stimulation (bottom) of synaptic transmission. **G.** Summary data of IPSC suppression by DAMGO and SNC162, comparing electrical to optogenetic stimulation (data are replotted from Figure 1D and 1I).

**Supporting Figure S2 (refers to Figure 4).**
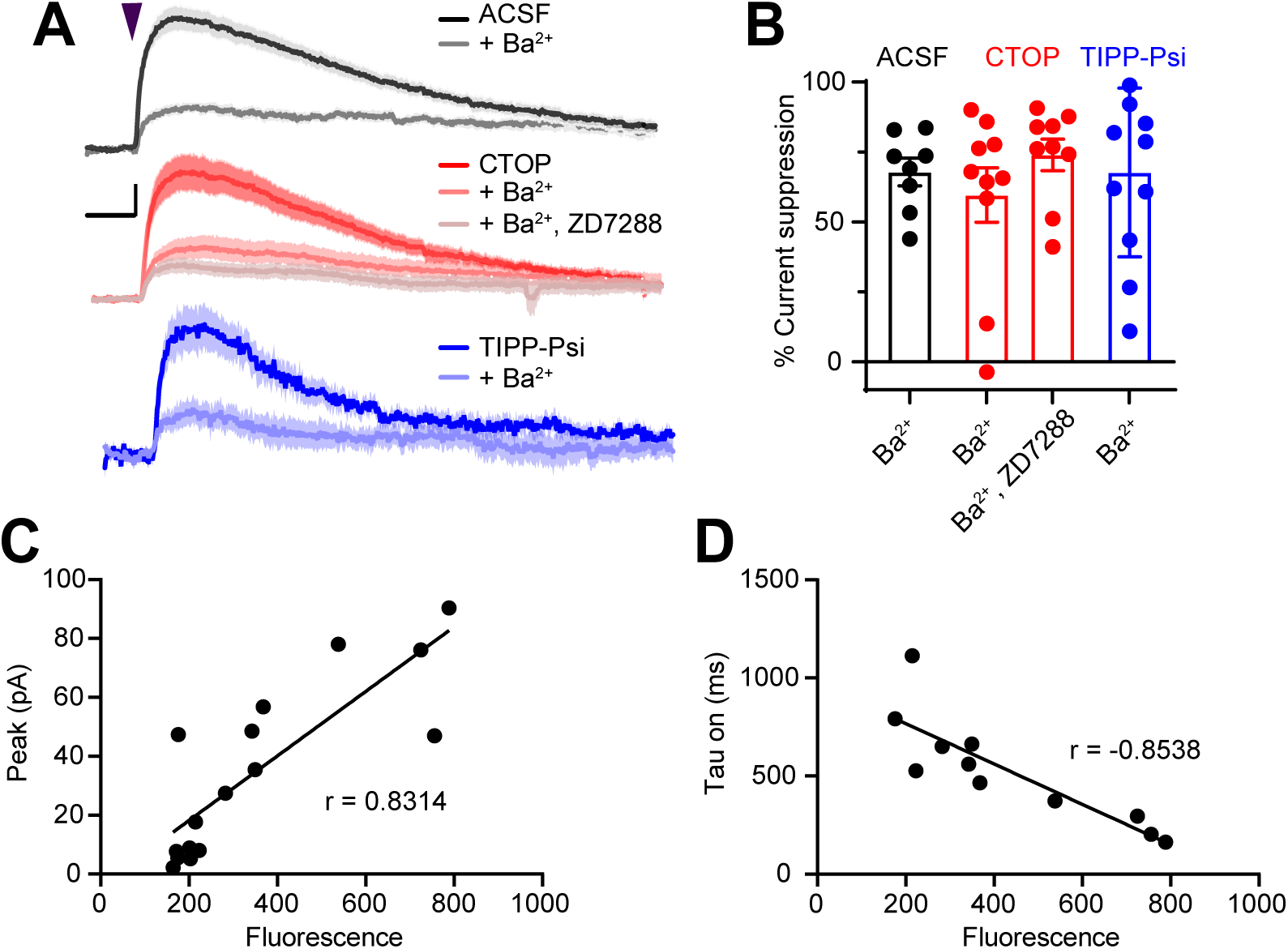
Sensitivity of somatodendritic currents to the GIRK blocker Ba^2+^ and mu opioid receptor expression level. **A.** Average outward currents evoked by photoactivation of CYLE with a 84 mW light flash in the absence (black, ACSF, n = 13 cells from 7 mice) and presence of mu and delta opioid receptor antagonists (red, CTOP, n = 14 cells from 10 mice; TIPP-Psi, n = 13 cells from 9 mice), as well as Ba^2+^ (1 mM) (gray, ACSF + Ba^2+^, n = 8 cells from 2 mice; light red, CTOP + Ba^2+^, n = 10 cells from 4 mice; light blue, TIPP-Psi + Ba^2+^, n = 10 cells from 4 mice), and the HCN blocker ZD7288 (lightest red, CTOP + Ba^2+^, ZD7288, n = 9 cells from 3 mice), as indicated. Scale bar: x = 5 sec, y = 25% of current without Ba^2+^ or ZD7288. **B.** Summary of the percentage of the average peak current amplitude that is blocked in each condition shown in **A**. **C.** Peak amplitude of the MOR-current in PV-Cre neurons expressing mCh-2A-hMOR vs red fluorescence in the recorded cell, as well as the Pearson’s correlation coefficient. **D.** Time constant of MOR-current activation in PV-Cre neurons expressing mCh-2A-hMOR vs red fluorescence in the recorded cell, as well as the Pearson’s correlation coefficient.

## Notes

### Competing Interest Statement

The authors have declared no competing interest.

